# Systematic Normalization with Multiple Housekeeping Genes for the Discovery of Genetic Dependencies in Cancer

**DOI:** 10.1101/2020.01.29.925651

**Authors:** Oliver Bonham-Carter, Yee Mon Thu

**Affiliations:** Allegheny College, Dept. of Computer Science, Meadville, PA; Allegheny College, Dept. of Biology, Meadville, PA

**Keywords:** cancer, gene expression, genome instability genes, housekeeping genes, normalization analysis

## Abstract

Cancer results from complex interactions between genes that are misregulated. Although our understanding of the contribution of single genes to cancer is expansive, the interplay between genes in the context of this devastating disease remains to be understood. Using the Genomic Data Commons Data Portal through National Cancer Institute, we randomly selected ten data sets of breast cancer gene expression, acquired by RNA sequencing to be subjected to a computational method for the exploration of genetic interactions at a large scale. We focused on genes that suppress genome instability (GIS genes) since function or expression of these genes is often altered in cancer.

In this paper, we show how to discover pairs of genes whose expressions demonstrate patterns of correlation. To ensure an inter-comparison across data sets, we tested statistical normalization approaches derived from the expression of randomly selected single housekeeping genes, or from the average of three. In addition, we systematically selected ten housekeeping genes for the purpose of normalization. Using normalized expression data, we determined *R*^2^ values from linear models for all possible pairs of GIS genes and presented our results using heatmaps.

Despite the heterogeneity of data, we observed that multiple gene normalization revealed more consistent correlations between pairs of genes, compared to using single gene expressions. We also noted that multiple gene normalization using ten genes outperformed normalization using three randomly selected genes. Since this study uses gene expression data from cancer tissues and begins to address the reproducibility of correlation between two genes, it complements other efforts to identify gene pairs that co-express in cancer cell lines. In the future, we plan to define consistent genetic correlations by using gene expression data derived from different types of cancer and multiple gene normalization.

**CCS CONCEPTS:**

- Applied computing → *Computational biology*.

**ACM Reference Format:** Oliver Bonham-Carter and Yee Mon Thu. 2019. Systematic Normalization with Multiple Housekeeping Genes for the Discovery of Genetic Dependencies in Cancer. In *Niagara Falls, New York.* ACM, New York, NY, USA, 10 pages. https://doi.org/10.1145/nnnnnnn.nnnnnnn

## 1 INTRODUCTION

The human genome sustains an incredible level of stability given a myriad of extrinsic and intrinsic insults that create alterations and potentially compromise the stability. Instability at a small-scale level could be beneficial since this is the source of genetic diversity. However, genome instability, at a large-scale level, may inadvertently introduce changes within the genome, which could result in deleterious outcomes. In multi-cellular organisms, genome instability is a precursor for genetic diseases such as cancer. Fortunately, multiple cellular mechanisms are in place to ensure that most forms of acquired damage do not perpetuate over multiple generations. These mechanisms can generally be categorized as pathways that promote repair, those that prevent inheritance of unintended alterations, and those that ensure cellular destruction, if the acquired damage is irreparable. Genes that are involved in these processes can collectively be designated as genome stability genes. Not surprisingly, many genes of genomic stability exhibit changes in expression or function in multiple cancers.

These changes provide a survival advantage for cancer cells since they foster genetic diversity within the population [1]. Nevertheless, how individual cancer cells thrive with the burden of a compromised genome is perplexing since multiple complex and redundant pathways exist to hinder propagation of the damaged genome. While healthy cells may undergo cell death or cell cycle arrest when genome integrity is compromised, cancer cells may evolve to prosper under the same conditions. These observations suggest that cancer cells, having acquired mechanisms to fend off the challenge of genome instability, have been selected during microevolution (i.e., during the tumor’s life cycle). Simultaneously, this implies that cancer cells may become dependent on these pathways to survive genome instability.

Identifying these pathways will reveal vulnerabilities of cancer. This approach has been successfully implemented in the design of targeted cancer therapies. For instance, shown in Figure 1-A, breast cancer cells with a mutation in DNA repair genes, *BRCA1* or *BRCA2*, are more sensitive to the inhibition of another repair pathway, mediated by the protein product of *PARP1*, than healthy cells without the mutation [7]. This sensitivity is as a result of *BRCA1/2* mutant cells’ dependency on the function of *PARP1*. Identifying additional vulnerabilities of cancer cells, with mutations that compromise genome stability, will help to encourage novel therapeutic approaches.

**Figure 1:**
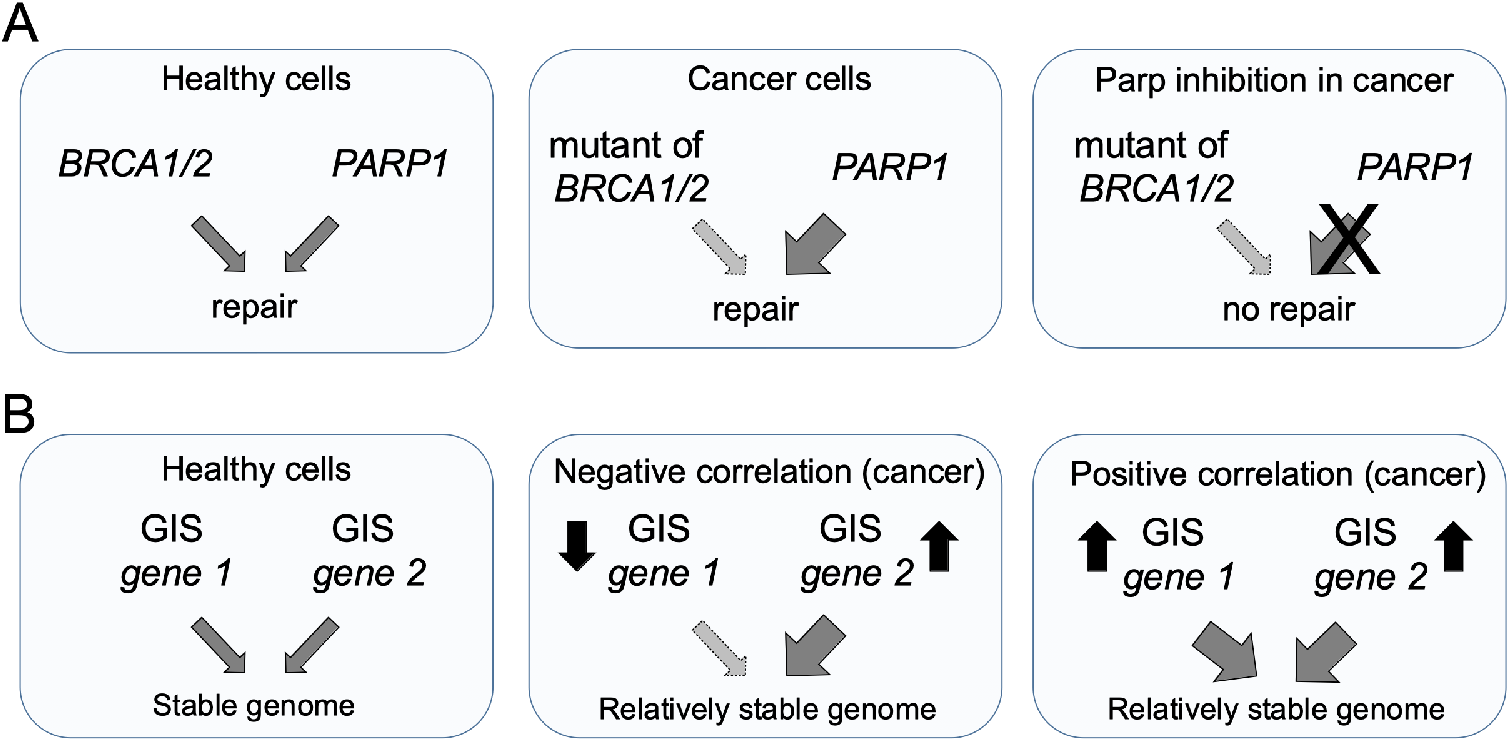
Genetic interactions between genes responsible for DNA repair or genome stability. **(A)** Cancer cells with mutated *BRCA1/2* genes become more dependent on the function of the *PARP1* protein product. As a result, they are more vulnerable to inhibition of *PARP1* compared to healthy cells, in which the protein products of *BRCA1/2* genes are functional [7]. **(B)** A working model to illustrate the genetic dependencies in cancer. In cancer cells, physiological expression levels of GIS genes may be altered (indicated by the up- or down-arrow). A change in the expression of one gene may prompt the dependency of another GIS gene, resulting in a proportional increased or decreased expression of the latter. The nature of this relationship may be manifested in a positive or negative correlation between the expressions of two genes.

To identify cancer cell vulnerabilities, we decided to examine the correlation of gene expressions using a large-scale approach. We reasoned that expression of two genes would positively or negatively correlate if the change in the expression of one gene necessitated the function of another, shown in Figure 1-B.

Due to the immense complexity of the study of expression in the genome, many computational methods, reviewed in [16] are necessary. The use of computational methods, where complex and voluminous amounts of data is involved, is seemingly the only approach for conclusive study, as discussed in [11] (differential analysis), [5] (signature study), [10] (expression profiling), [17] (cancer-specific correlations) and other studies mentioned in [3]. For our own study of diverse pathways, there was no substitution to automated methods. Therefore, to gain insight into gene expression correlation in cancer, we elected to include computational methods from Bioinformatics.

We used gene expression data from the Genomic Data Commons Data Portal (GDC) through National Cancer Institute (NCI). From these data sets, we selected a subset of genes that suppress genome instability (GIS genes), as categorized in the study by Putnam *et al.* [13]. These genes were regarded in the author’s article as GIS genes since perturbation in the function of these genes lead to small-scale or large-scale changes within the genome.

In this study, we use FPKM datasets to show how to determine the existence of a positive or negative correlation between the expression of two given GIS genes in cancer, as shown in Figure 1-B. These correlations can reveal if two GIS genes coordinates or if alteration in the expression of one GIS gene increases dependency of cancer cells on another GIS gene, as seen in Figure 1-B. Co-expression between pairs of genes has been studied in cancer cell lines [9]. However, to our knowledge, a similar type of analysis has not been performed using gene expression data from cancer tissues. Furthermore, it would be beneficial to determine the reproducibility of these correlations. Here, we report a preliminary data on co-expression of pairs of GIS genes using breast cancer gene expression data. We observed some consistent correlations between certain pairs although different groups of housekeeping genes were used for normalization. In the future, our goal is to expand the number of data sets in order to identify pairs of genes that exhibit consistent co-expression.

## 2 METHODS

The *Transcriptome Profiling* data for this project was downloaded from the Genomic Data Commons (GDC) Data Portal, (https://portal.gdc.cancer.gov/) available from the National Cancer Institute (NCI). The data that we selected for our work was arbitrarily chosen for its association with Breast Invasive Carcinoma and its affiliation with the GDC’s “TCGA-BRCA” listing, as specified by the GDC website. In Table 1, we display the ID’s and file names of the individual sets used for the current work. Each of our chosen sets were from GDC’s *Transcriptome Profiling* category, of type *Gene Expression Quantification* and RNA-Seq experimental strategy.

**Table 1:**
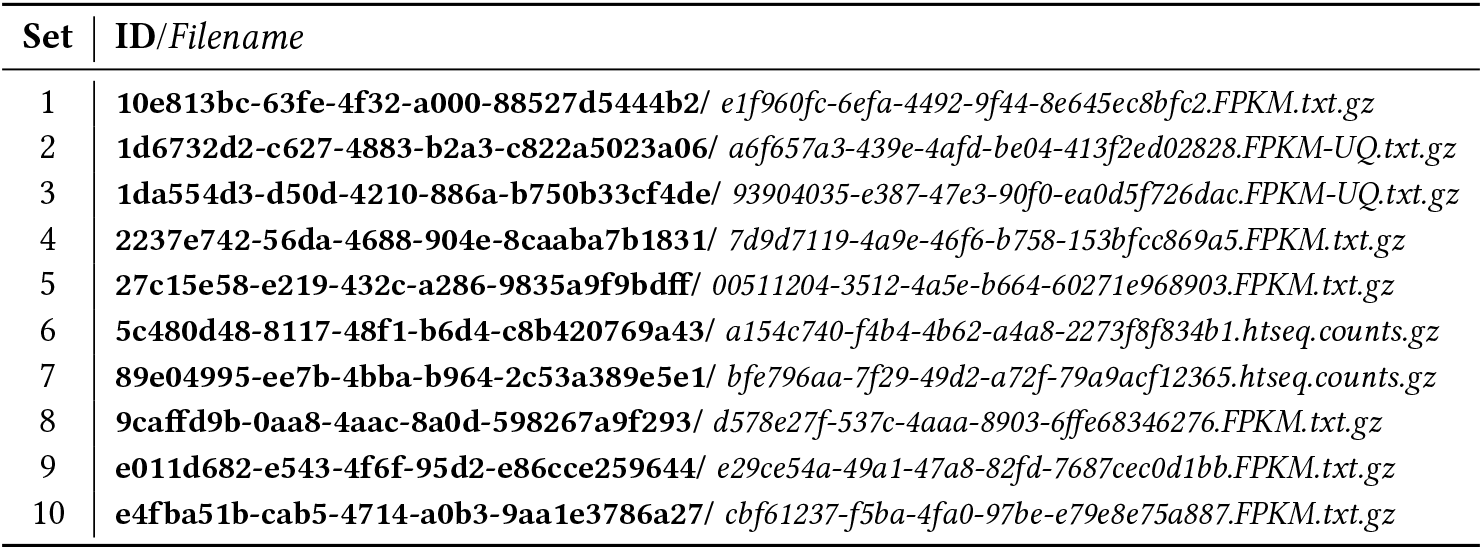
The gene expression data downloaded from the GDC Data Portal available from NCI. The ID and the data set make up the directory and filename of the data set (i.e., `ID/data set.ext`.) In this work, each set is labeled by the index *d*, for 1 ≤ *d* ≤ 10.

All code (available upon request) and data was handled using RStudio [15] for R-Statistics [14]. We defined our *superset* of gene expressions to imply the complete listing of Ensemble IDs originating from each file in our data set of Table 1. Each file contained exactly the same listing of Ensemble IDs and their associated gene expression values, which vary from data set to data set.

From the human expression data described in Table 1, we defined a subset of 273 GIS genes (i.e., genes that are responsible for maintaining integrity of the genome), which were the focus for our study. GIS genes were obtained from the study by Putnam *et al.* [13]. This subset was important for two general reasons, (1) expression of these genes directly addressed our research question mentioned in Introduction, and (2) we determined that there was too much gene diversity to cover in the total data (60484 genes total) to permit a concentrated study.

Due to the inherent noise and other sources of error associated with the quantification and collection of expression data, normalization is necessary to allow for studies of comparison. The overarching goal of our study was to determine normalizing factors that make it possible for us to discover pairs of GIS genes which show consistent correlation. To this end, we studied single and multiple gene expressions to be used to normalize data, as discussed below.

### 2.1 Single Expression Normalization

Our first step was to study the possibilities of normalization using the expression of single genes as inputs in our calculations. For this, we arbitrarily selected the following three housekeeping genes; *TUBB* (ENSG00000196230.11), *TUBA1A* (ENSG00000167552.12) and *GAPDH* (ENSG00000111640.13).

In each of our data sets, the gene expression values for selected genes were obtained and applied to the normalization of the set (i.e., those featured in Table 1) from which the particular expression values were obtained. The names of our normalizing variables follow the convention set-out in Table 2. In this table, one will note that the expression of a housekeeping gene (*TUBB*, *TUBA1A* or *GAPDH*) in each data set is used as a normalizing factor for that data set. In other words, for single expression normalization, each data set has three normalizing factors, that are unique to that set.

**Table 2:**
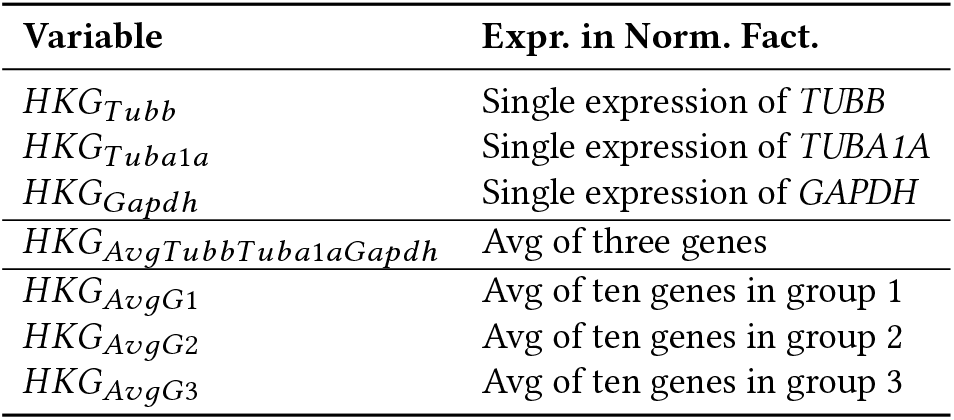
The legend of variable names of the normalizing factors that were used to normalize the gene expressions. For each set, these variables were created to be used to normalize all expressions of the set.

For example, the normalizing factor that was created for each data set using the *TUBB* gene expression, can be written, *HKG_Tubb_* and similarly for the other genes, *TUBA1A* (*HKG*_*Tuba*1*a*_) and *GAPDH* (*HKG*_*Gapdh*_). Each data set featured in Table 1 has its own specific *HKG*_*Tubb*_, *HKG*_*Tuba*1*a*_ and *HKG*_*Gapdh*_ values.

Using these values, we calculated a normalizing expression for each GIS gene (total N = 272) for each data set in Table 1. The natural log of the normalized values, denoted by, *y_n_*, of all GIS genes (*gene*_*n*_ for 1 ≤ *n* ≤ *N*) were calculated from expression values (denoted by *x*_*n*_) using the normalizing factors featured in Table 2. For example, the equation for normalized expressions in a set using *TUBB* as a normalizing factor (*HKG*_*Tubb*_) was the following.

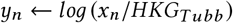

With log-transformation, we were able to achieve normal distribution of values, that may otherwise be skewed [2].

### 2.2 Multiple Expression Normalization

#### Three genes

In an effort to diversify the normalizing factors for each data set featured in Table 1, we used the average of the expressions of the three genes described above (i.e., *TUBB*, *TUBA1A* or *GAPDH*) instead of expression of individual genes. For example, described in Figure 2, the natural log of the normalized value (denoted by *y_n_*) for each GIS gene (*gene_n_* where 1 ≤ *n* ≤ *N*) was obtained in each data set by the following equation.

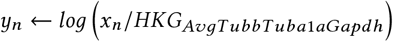

**Figure 2:**
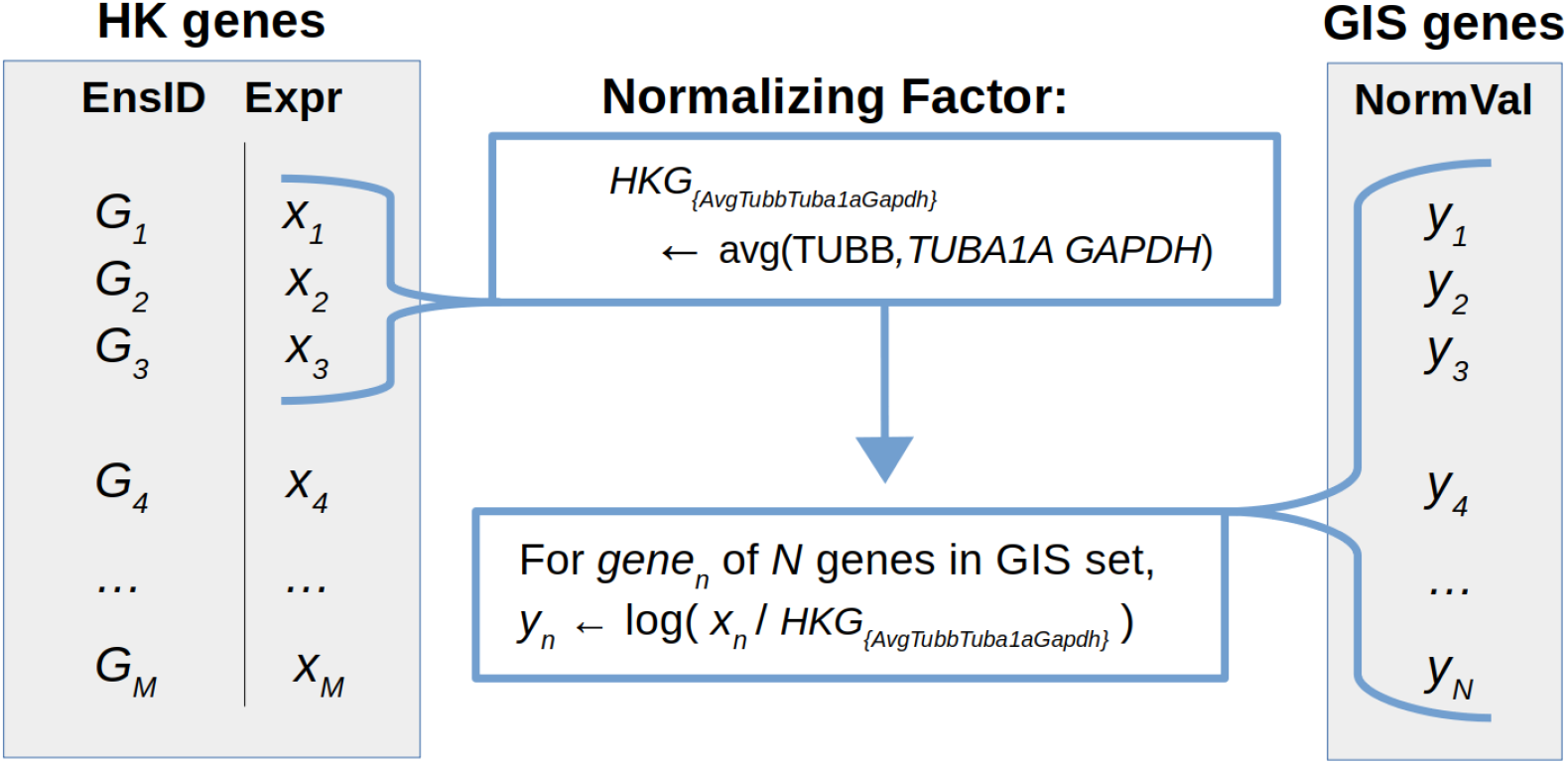
Each expression value was normalized according to an average derived from selected gene expression values in the same data set. This average was calculated in each data set using the selected Ensemble IDs.

#### Ten genes

For this approach, we used the average of ten gene expressions to obtain normalizing factors for each set shown in Table 1. We speculated, that the average of ten genes, chosen for their functional attributes, would create a normalizing factor that would better represent the diversity of housekeeping genes.

As part of this approach, we first identified a list of approximately 3800 housekeeping genes, whose expressions are relatively uniform across multiple tissues [6]. This is an important criteria since we intend to use this normalization method for comparison of gene expression derived from cancers of different tissue of origins. Housekeeping genes from this list were subject to gene ontology analysis (http://geneontology.org/) so that we could systematically categorize them based on cellular components to which they belonged.

After sorting according to fold enrichment values, the top ten non-redundant cellular components were chosen and one representative gene from each functional category was randomly selected to build a list of housekeeping genes. The same process was repeated to generate two additional lists of housekeeping genes. We identify the selected housekeeping genes and their ontologies in Tables 4, 5, and 6. We speculated that we would get consistent results if ten genes that we selected correctly represented the diversity of house-values, as shown keeping genes. The average expression of ten genes in each list was determined, denoted by, *HKG*_*AvgG*1_, *HKG*_*AvgG*2_ and *HKG*_*AvgG*3_. These three averages were determined to obtain three normalizing factors unique for each set in Table 1.

**Table 3:** The variance and mean of values described in R^2^ heatmaps across normalizing factors. The heatmaps were designed to illustrate trends in the numbers of gene-gene correlations according to their elevated R^2^ values. Too many high mean values (as noted in the single gene) are biologically suspect and we noted that when ten genes were used to create the normalizing factors, there were fewer gene-gene correlations (lower means). These results are in-keeping with the natural high-complexity to make such gene-gene interactions possible in biology [4].

| Normalizing Factor | Mean | Variance |
| --- | --- | --- |
| $HKG_{TUBB}$ | 0.28617 | 0.06526 |
| $HKG_{Tuba1a}$ | 0.71638 | 0.08067 |
| $HKG_{GAPDH}$ | 0.55054 | 0.12963 |
| $HKG_{AvgTubbTuba1aGapdh}$ | 0.26050 | 0.05792 |
| $HKG_{AvgG1}$ | 0.22230 | 0.05274 |
| $HKG_{AvgG2}$ | 0.19935 | 0.04645 |
| $HKG_{AvgG3}$ | 0.20610 | 0.04732 |

**Table 4:** The first set of housekeeping genes for multiple expression normalization. In the group, the expression values of all ten were averaged in each set *d* to determine its normalizing factor. Here, we provide the ontology group, Uniprot ID, *human* gene ID and Ensemble IDs.

| Ontology | UnitProtKB | Gene ID | EnsNum |
| --- | --- | --- | --- |
| Proteasome | Q9Y5K5 | *UCHL5* | ENSG00000116750 |
| Ribosome | Q96EL2 | *MRPS24* | ENSG00000062582 |
| ER | Q9P0I2 | *EMC3* | ENSG00000125037 |
| ER targeting | O76094 | *SRP72* | ENSG00000174780 |
| splicing | O15234 | *CASC3* | ENSG00000108349 |
| Prp19 complex (splicing) | Q9BZJ0 | *CRNKL1* | ENSG00000101343 |
| splicing snRNP complex | P62310 | *LSM3* | ENSG00000170860 |
| Proteasome | P62195 | *PSMC5* | ENSG00000087191 |
| Methylome | Q9BQA1 | *WDR77* | ENSG00000116455 |
| Translation | Q9UBQ5 | *EIF3K* | ENSG00000178982 |

**Table 5:** The second set of housekeeping genes for multiple expression normalization.

| Ontology | UnitProtKB | Gene ID | EnsNum |
| --- | --- | --- | --- |
| Proteasome | Q9NRR5 | *UBQLN4* | ENSG00000160803 |
| Ribosome | Q9H2W6 | *MRPL46* | ENSG00000259494 |
| ER | Q8N766 | *EMC1* | ENSG00000127463 |
| ER targeting | P37108 | *SRP14* | ENSG00000140319 |
| splicing | P38919 | *EIF4A3* | ENSG00000141543 |
| Prp19 complex (splicing) | Q99459 | *CDC5L* | ENSG00000096401 |
| splicing snRNP complex | Q53GS9 | *USP39* | ENSG00000168883 |
| Proteasome | Q06323 | *PSME1* | ENSG00000092010 |
| Methylome | O14744 | *PRMT5* | ENSG00000100462 |
| Translation | Q7Z478 | *DHX29* | ENSG00000067248 |

**Table 6:** The third set of housekeeping genes for multiple expression normalization.

| Ontology | UnitProtKB | Gene ID | EnsNum |
| --- | --- | --- | --- |
| Proteasome | P14735 | *IDE* | ENSG00000119912.14 |
| Ribosome | Q96BP2 | *CHCHD1* | ENSG00000172586.7 |
| ER | Q9BV81 | *EMC6* | ENSG00000127774.6 |
| ER targeting | P49458 | *SRP9* | ENSG00000143742.11 |
| splicing | Q9HCG8 | *CWC22* | ENSG00000163510.12 |
| Prp19 complex (splicing) | P11142 | *HSPA8* | ENSG00000109971.12 |
| splicing snRNP complex | O43447 | *PPIH* | ENSG00000171960.9 |
| Proteasome | Q15008 | *PSMD6* | ENSG00000163636.9 |
| Methylome | Q99873 | *PRMT1* | ENSG00000126457.19 |
| Translation | Q7L2H7 | *EIF3M* | ENSG00000149100.11 |

Using *HKG_AvgG_*_1_ unique for each data set, the natural log of normalized values (denoted by *y_n_*) for each GIS *gene_n_* where 1 ≤ *n* ≤ *N*) was obtained by the following equation.

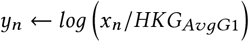

The same procedure was performed using corresponding *HKG*_*AvgG*2_ and *HKG*_*AvgG*3_.

### 2.3 Regression models

When the gene expressions of a data set are normalized by a single or multiple expression normalizing factor, we determined correlation of expression between all possible pairs of genes within the GIS list. Correlation is represented by R^2^ values, generated from a test where the normalized expression values of GIS genes underwent an *all-against-all* linear regression test. To visualize the whole data set, we used heatmaps in which R^2^ values are represented as a gradient of colors. The R^2^ statistical metric is a measurement of proximity of data points to the fitted regression line. Also known as a “coefficient of determination”, this statistic describes the percentage of the response variable variation, as explained by a linear model and is on a scale that ranges from 0 to 100 percent. The values in our heatmaps range between 0 and 1, in keeping with the lower- and upper-bounds, respectively, of their R^2^ values. The mean and variance values were collected for each normalizing factor and are shown in Table 3.

In Figure 3, we describe the steps to create heatmaps from the R^2^ values of linear models between pairs of genes. The generation of heatmaps from normalizations using normalizing factor values (i.e., single or multiple gene expressions) was similar across all prepared figures. Our heatmap graphics were prepared using the Plotly library in R [8].

**Figure 3:**
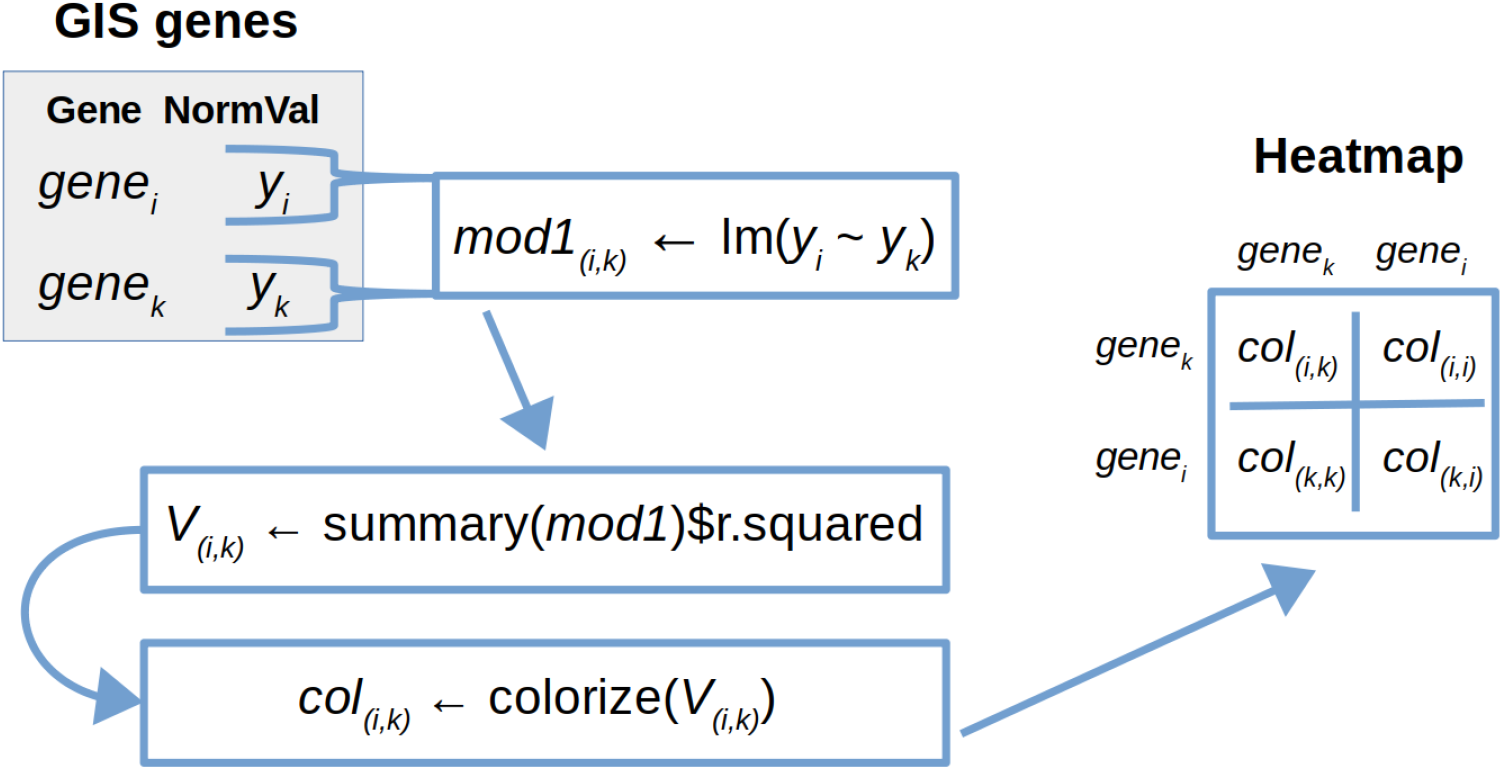
The R^2^ values from linear models where the normalized values of one gene is regressed over another are collected. We note that *gene_i_* and *gene_k_* represent all normalized values obtained across the data sets of Table 1. The heatmaps are generated from R^2^ values from linear regression models. The values have been colorized according to their placement in the natural range of R^2^ (i.e., [0 ≤ *R*^2^≤ 1].)

## 3 RESULTS AND DISCUSSION

In this study, we aimed to develop a normalization approach that allows us to identify consistent correlation between expression of two GIS genes. We used gene expression data available through NCI GDC data portal (https://portal.gdc.cancer.gov/). Ten gene expression data sets generated by RNA-sequencing of breast cancer tissues were randomly chosen for our data analysis. We employed different new normalization approaches to compare gene expression across multiple samples, as detailed in the Methods section.

When the *TUBB* expression was used for normalization, almost half of the gene pairs showed correlations, as shown in Figure 4. However, this level of codependency between genes is biologically improbable, suggesting that most of the data are false positives. When the expression values were normalized using another single gene expression value (*TUBA1A*), we noted that there were also many false positives in terms of R^2^ values showing high correlations, as shown in Figure 5. Furthermore, this result indicated a lack of consistency among single expression normalizations.

**Figure 4:**
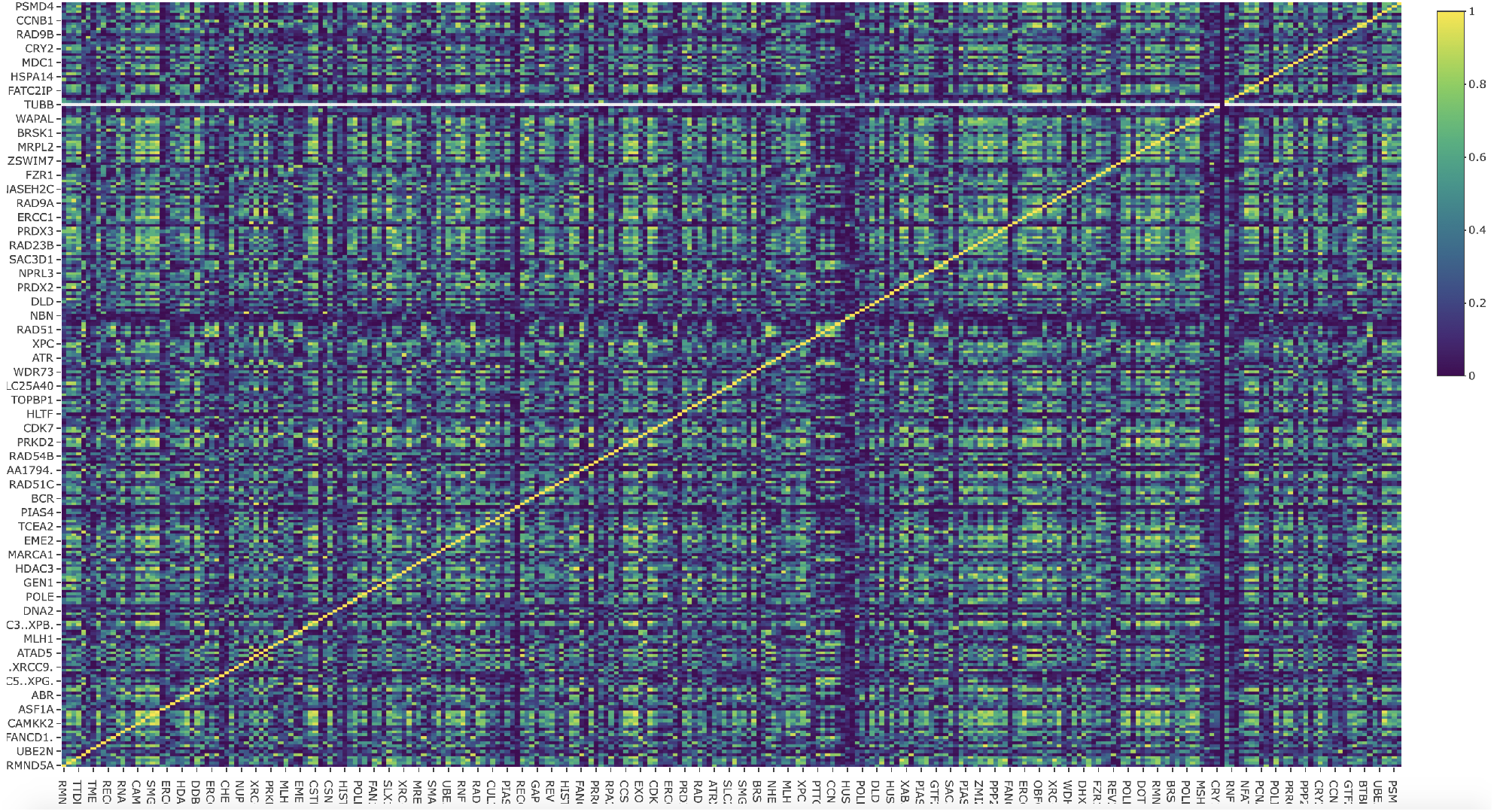
Heatmaps of R^**2**^ values, derived from the normalizing factor, *HKG_Tubb_*.

**Figure 5:**
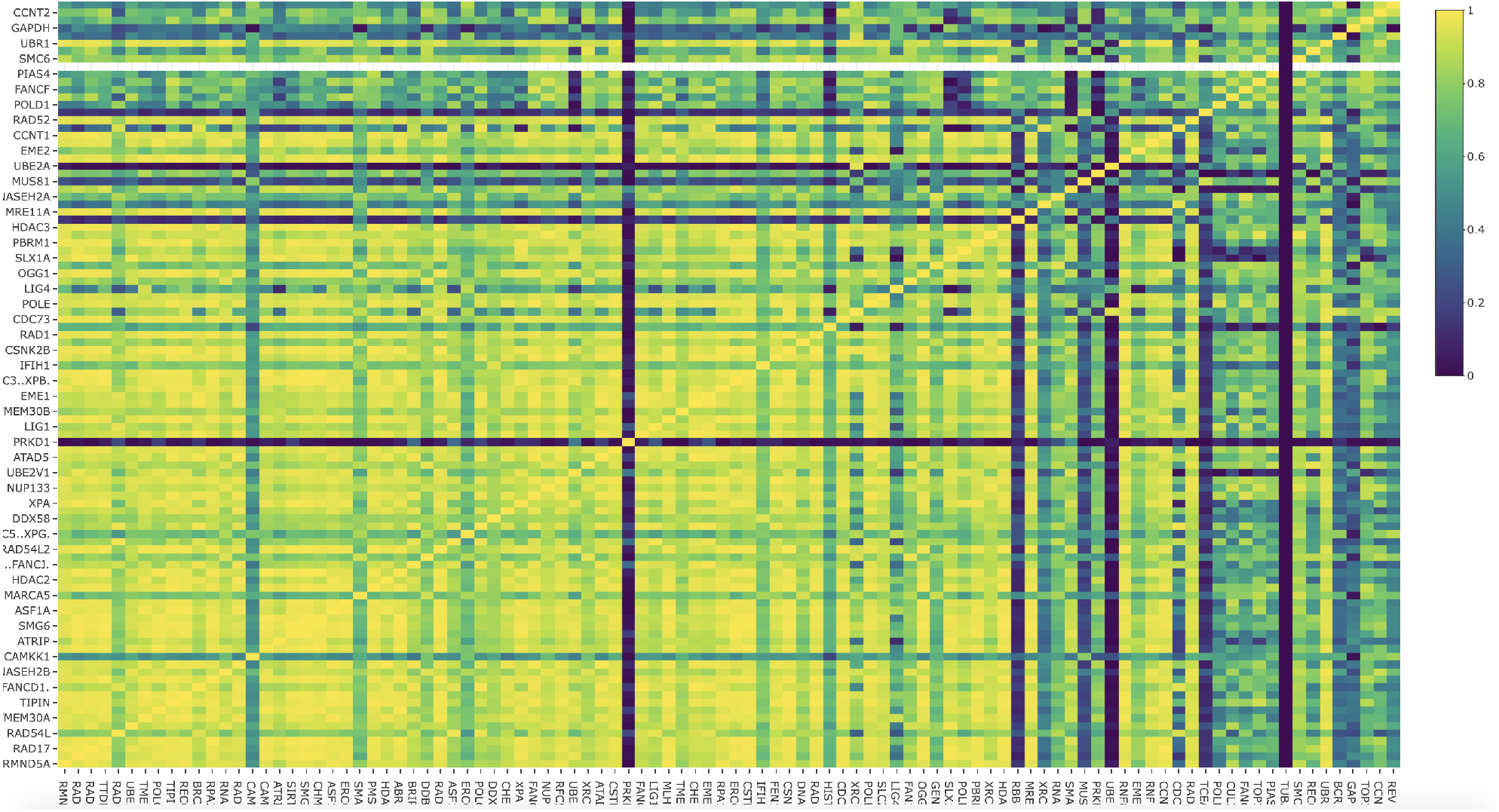
Heatmaps of R^2^ values, derived from the normalizing factor, *HKG_Tuba_*_1*a*_.

Normalizing with *HKG*_*AvgTubbTuba*1*aGapdh*_ slightly reduced the number of pairs with high linear correlation in Figure 6. This observation suggests that using only three housingkeeping genes is not sufficient to reduce false positives. In the gene normalization survey by Vandesompele *et al.* [20], ten house keeping genes were sufficient for normalization purposes, whereas, single genes selected for normalization often led to misleading results.

**Figure 6:**
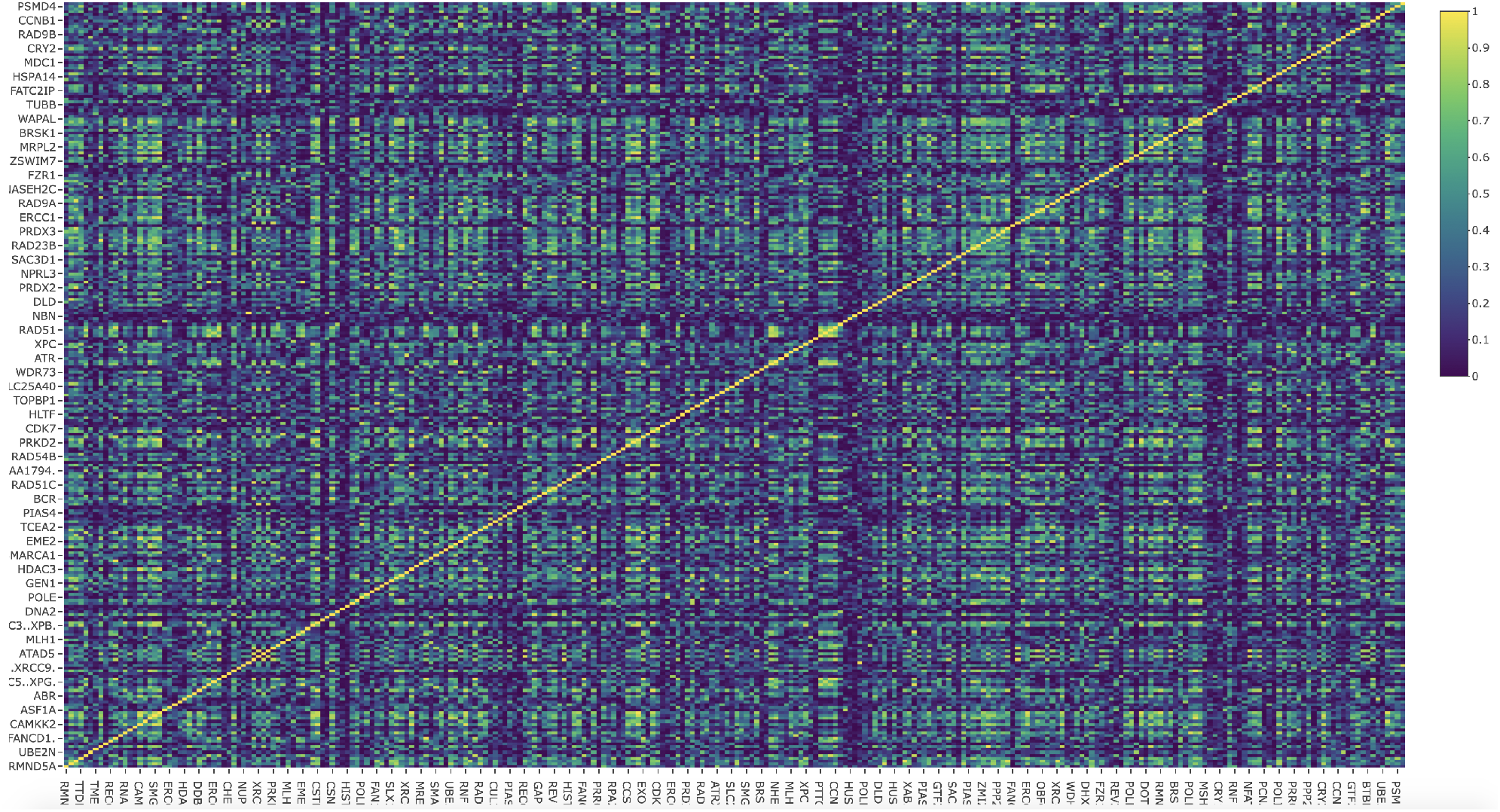
Heatmaps of R^2^ values, derived from the normalizing factor, *HKG_AvgTubbTuba_*_1*aGapdh*_.

We therefore applied the average expression of ten housekeeping genes for normalization, as described in the Methods of Section 2. Interestingly, the heatmap generated using *HKG*_*AvgG*1_, *HKG*_*AvgG*2_ or *HKG*_*AvgG*3_, of Figures 7, 8 and 9, respectively, shows a pattern similar to the counterpart generated by the *HKG_AvgTubbTuba_*_1*aGapdh*_ normalization, but was significantly different from the one generated by the *TUBB* normalization. Pairs with R^2^-values close to 1 were comparatively rare in the heatmaps of Figures 7, 8 and 9 for normalizing factors, *HKG*_*AvgG*1_, *HKG*_*AvgG*2_, and *HKG*_*AvgG*3_, respectively.

**Figure 7:**
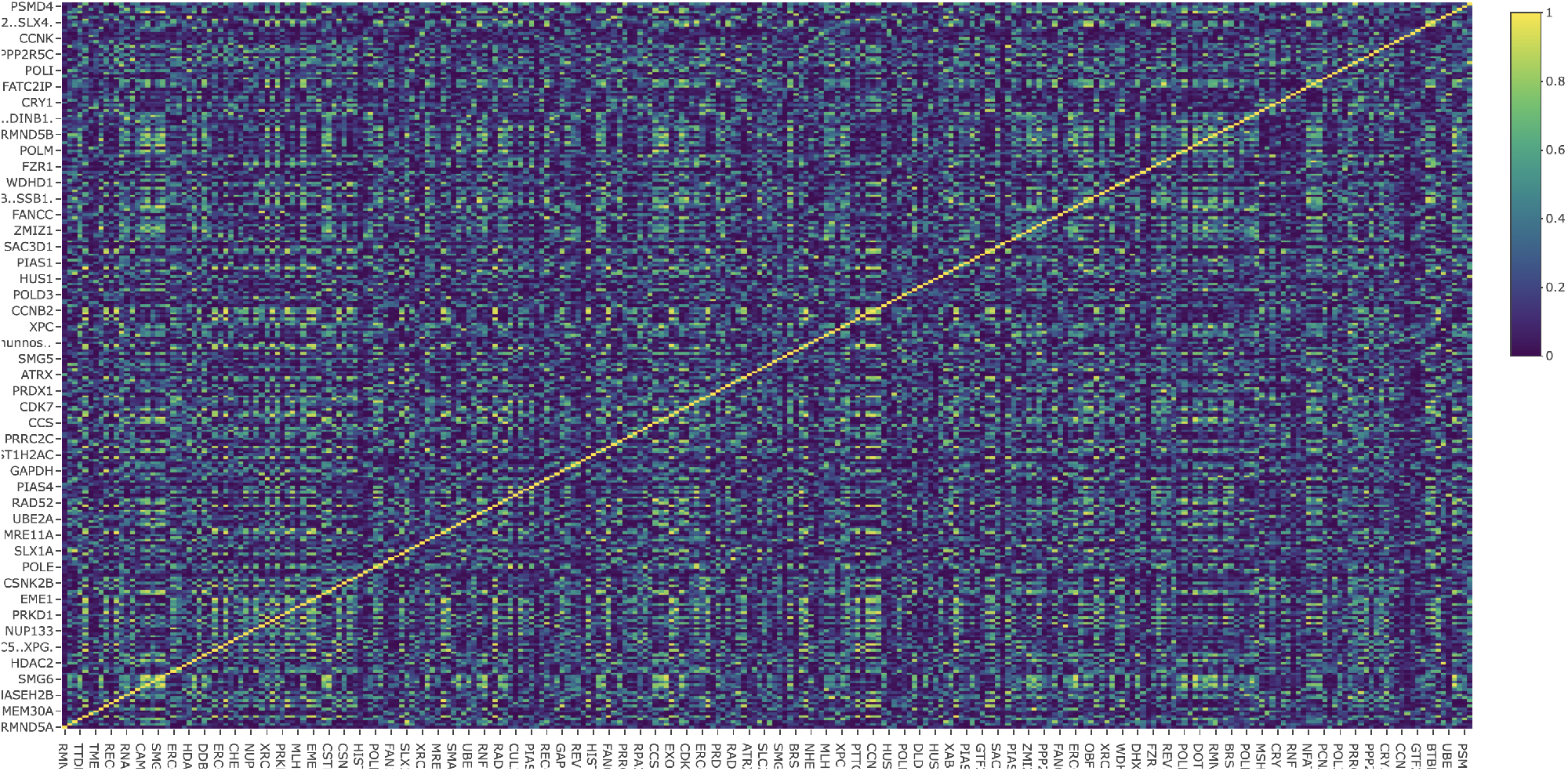
Heatmaps of R^2^ values, derived from the normalizing factor, *HKG*_*AvgG*1_.

**Figure 8:**
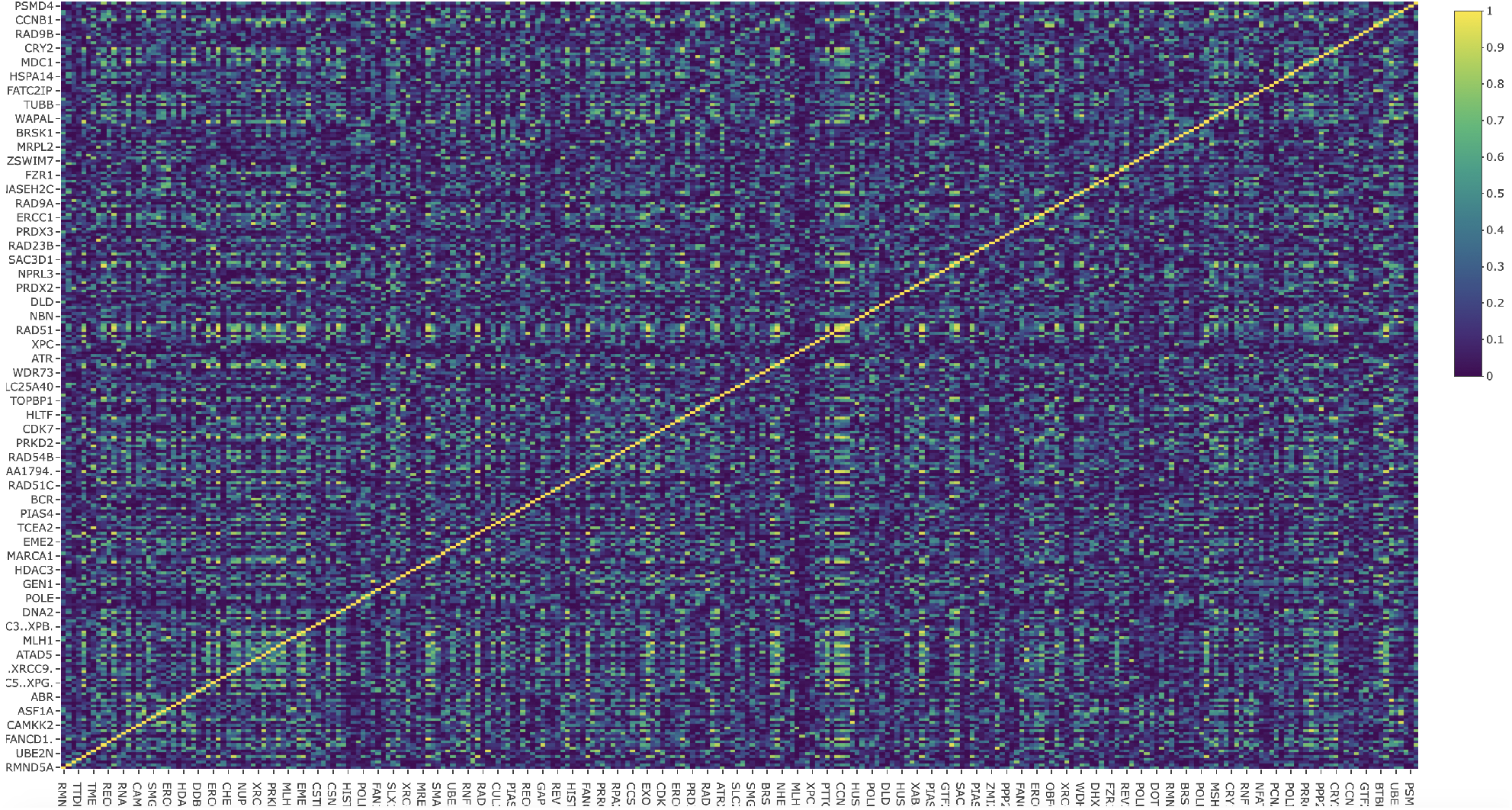
Heatmaps of R^2^ values, derived from the normalizing factor, *HKG*_*AvgG*2_.

**Figure 9:**
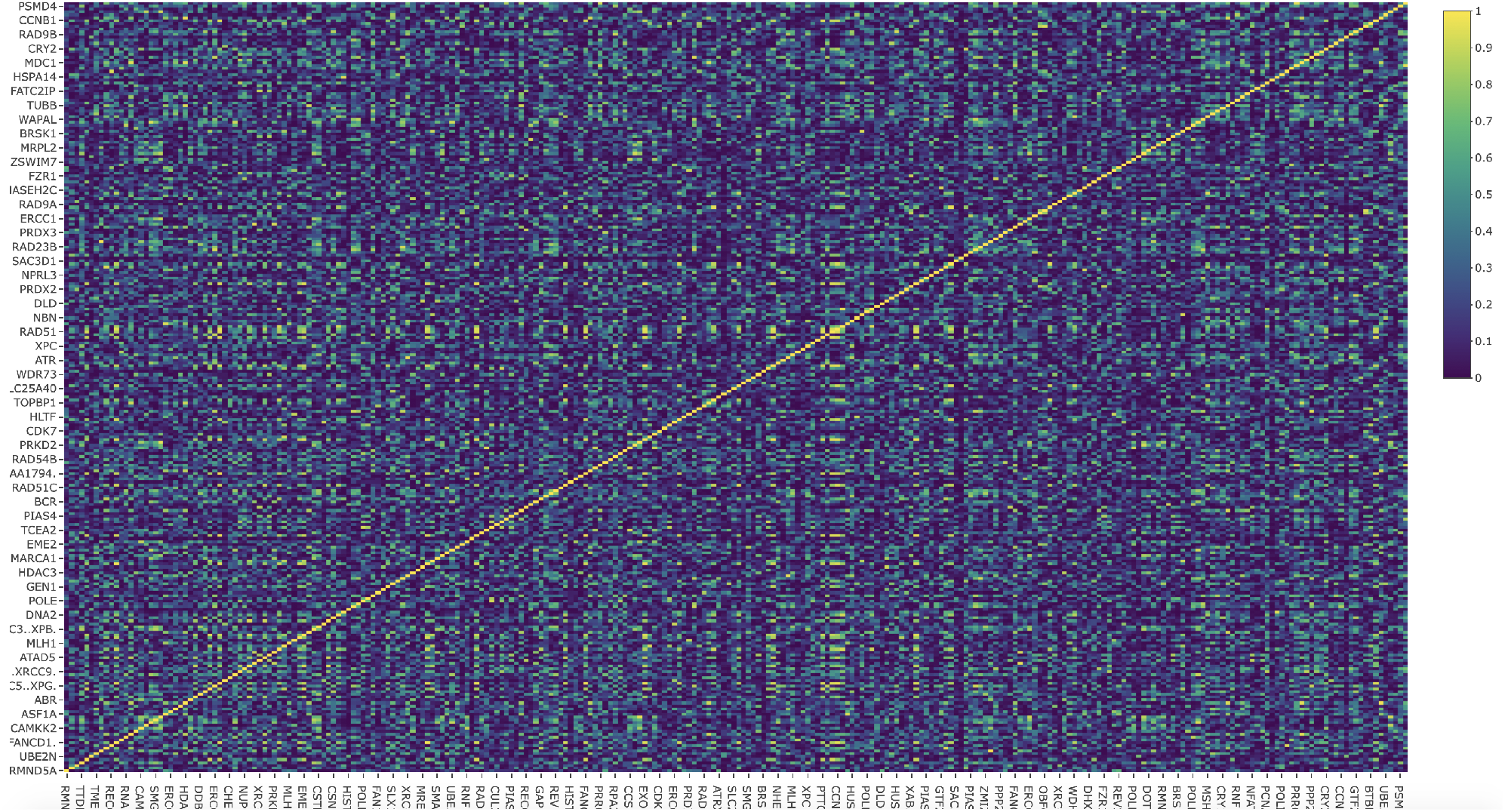
Heatmaps of R^2^ values, derived from the normalizing factor, *HKG*_*AvgG*3_.

Shown in Table 3, the variance and mean values were collected from each normalizing factor experiment. We noted that when the normalizing factors were larger, the gene-gene correlations appeared to be more reliable in spite of the natural high complexity of gene-gene interactions [4].

If these correlations are true positives, the pattern should be reproducible regardless of what normalizing factors were used. In support of this idea, the heatmaps generated by average of ten expression values appear similar to each other. In other words, correlation in expression is preserved for certain pairs of genes regardless of what normalizing factors are used, as shown in all heatmaps in Figures 7, 8 and 9. In future works, we intend to construct experiments with more statistical analysis across larger data sets to be able to determine more of the wide-spread consistencies concerning gene-gene correlations.

Linear correlation between two genes imply that alteration in expression of one gene necessitates a proportional response of the other, shown in Figure 1-B. The idea of genetic dependency stems from “genetic interaction,” a concept that has been long rooted in classical genetics. Large-sale genetic interactions have been determined with relative ease in simple eukaryotic model organisms, such as *Saccharomyces cerevisiae* [19].

In general, genetic interaction between two genes exist if the absence of both genes produces a phenotype, that is different from phenotypes resulting from deficiency of individual genes [12]. For instance, combined deficiency of two genes result in cell death, whereas deficiency of each gene permits cellular survival with reduced fitness. This scenario exemplifies a type of widely known genetic interaction and is succinctly referred to as “synthetic lethality.” This idea was first described by Bridges in 1922, and has been proposed to use for cancer therapy since 1997 by Hartwell and Friend [12].

Since then, the search for synthetic lethal interactions between genetic conditions specific to cancer has been ongoing. The genetic screen performed in cancer cells, using either CRISPR or RNAi, to understand which gene expression changes are necessary for cancer with a specific genetic background reveal multiple synthetic lethal interactions [18, 21].

In addition, multi-pronged approach such as DAISY has been developed to further solidify identification of synthetic lethal gene pairs [9]. Our study complements these efforts due to two main reasons. First, co-expression analysis described in DAISY was performed using data derived from cell lines, whereas our study uses gene expression data from cancer tissues. Second, our preliminary study begins to address the reproducibility of co-expression that may be universal or tissue specific. With a more comprehensive analysis, we anticipate that our study will offer insights into genetic dependencies within patient tumor tissues.

## 4 CONCLUSIONS

In this work, comprising an automated, comparative, *all-against-all*, gene study, we discovered GIS genes whose expression values positively or negatively correlated with the expressions of other genes. Our results were gathered utilizing computational methods for FPKM gene expression normalization, allowing us to regress each gene over all the others in the study to collect adjusted R^2^ values, from which we were able to detect potential correlations. Heatmaps were used to visualize the R^2^ values to enable us to spot correlations according to elevated values.

To accomplish this goal, we determined normalizing factors (i.e., derived values from house keeping genes to be used to normalize the gene expression values of GIS genes) that served for comparison purposes across the unique data sets chosen for our study. An important part of this comparison involved linear models where the normalized expression data of genes was regressed over that of all other genes. In our study, consistency represented a plausible correlation between GIS gene expressions that were normalized by normalizing factors. Consistent correlations, we reasoned, would show patterns which would be also found from the successful normalizing factors that would be derived from the expressions of other housekeeping genes.

### Single Expression Normalization

We found that the normalizing factors derived from single housekeeping genes did not provide generally consistent or meaningful results. For instance, when using single genes to create the normalizing factors the results of each normalizing factor were dissimilar. In addition, our results described too many biologically improbable correlations. We rejected these results as false positives. In total, since the single expression values were generally unable to achieve consistent results with each other, we found that this approach was not effective.

### Multiple Expression Normalization

We found that using the averaged expression values of multiple housekeeping genes was an effective approach to finding consistency across our data sets. When we used the average of three genes to normalize expression data, we found less cases that we estimated were biologically improbable. However, on close inspection of the results, we still noted that there were a large number of likely false positives. We noticed a pattern emerge; the more genes we used to create normalizing factors, then the more consistent the results were according to correlations.

Our preliminary method enabled us to identify the co-expression of gene pairs in breast cancer tissues. This technique allows for reproducibility across data sets and to compare approaches involving diverse normalization factors when detecting correlation patterns.

In future studies, we will derive normalizing factors from other collections of ten (or more) genes to test for consistency and biologically relevant correlations in the data of other categories of cancer. For instance, we intend to use a similar approach to analyze larger number of data sets derived from different tumor types, in addition to breast cancer tissue. We will compare correlation data within and across tissue types to determine if the normalizing factors derived from ten housekeeping genes are suitable for both means of comparison. This analysis may also reveal correlation of gene expression that are unique to certain types of tissues. Using multiple expression normalization, we hope to identify pairs of genes which show consistent correlation regardless of normalizing factors or data sets.

